# Domain-invariant features for mechanism of action prediction in a multi-cell-line drug screen

**DOI:** 10.1101/656025

**Authors:** Joseph C. Boyd, Alice Pinheiro, Elaine Del Nery, Fabien Reyal, Thomas Walter

## Abstract

High Content Screening is an important tool in drug discovery and characterisation. Often, high content drug screens are performed on one single cell line. Yet, a single cell line cannot be thought of as a perfect disease model. Many diseases feature an important molecular heterogeneity. Consequently, a drug may be effective against one molecular subtype of a disease, but less so against another. To characterise drugs with respect to their effect not only on one cell line but on a panel of cell lines is therefore a promising strategy to streamline the drug discovery process. The contribution of this paper is twofold. First, we investigate whether we can predict drug mechanism of action (MOA) at the molecular level without optimisation of the MOA classes to the screen specificities. To this end, we benchmark a set of algorithms within a conventional pipeline, and evaluate their MOA prediction performance according to a statistically rigorous framework. Second, we extend this conventional pipeline to the simultaneous analysis of multiple cell lines, each manifesting potentially different morphological baselines. For this, we propose multitask autoencoders, including a domain-adaptive model used to construct domain-invariant feature representations across cell lines. We apply these methods to a pilot screen of two triple negative breast cancer cell lines as models for two different molecular subtypes of the disease.

## 1 Introduction

High content screening (HCS) is a powerful tool for identifying potential drugs effective against a particular disease. A high content drug screen corresponds to a series of imaging experiments under controlled conditions, where a cell line representative of some disease is exposed to a large panel of drugs. For each drug, one obtains a set of images informative of its phenotypic effect and hence on the biological pathways undergoing perturbation. Various advances in microscopy automation and image analysis have pushed HCS to the early *hit-to-lead* stages of the drug discovery process [11].

The discovery of new drugs may be guided by a reference set of drugs of known mechanism of action (MOA). The MOA of a drug is the particular cellular pathway it perturbs to achieve its effect. Through application of image analysis, one may attempt to infer the MOA of an unknown drug from HCS image data. Note that MOA can be defined at different levels and with different degrees of specificity: MOA might concern the exact protein that is targeted (e.g. AURKA inhibition), or a specific effect on cellular components (e.g. stabilisation of microtubuli) or perturbation of a more general cellular pathway (e.g. DNA repair). HCS is usually optimised with respect to particular pathways by the choice of the fluorescent markers and readouts [27]. Consequently, MOA prediction might be reasonably straightforward if MOA classes are chosen in accordance with the phenotypic readout [19], but it is challenging in general, in particular if we aim at predicting specific MOAs the assay has not been optimised for.

A second difficulty concerns the cellular model that is used. As a proxy for diseased cells, a cell line cannot be thought of as a perfect model. Many diseases feature a significant molecular heterogeneity. Consequently, a drug may be effective against one molecular subtype of a disease, but less so against another. Furthermore, immortalised cell lines may diverge over time due to genetic drift. For example, HeLa, the quintessential cell line, is famously the cause of great scientific confusion due to difficulties in cell line identification [13] and significant molecular and phenotypic variability [18]. To characterise drugs with respect to their effect not only on one cell line but on a consensus of several is therefore a promising strategy to streamline the drug discovery process. Nevertheless, this is not an easy task in morphological screening, as different cell lines usually have distinct archetypal morphologies even prior to perturbation. It is therefore conceptually difficult to characterise and compare drug effects across cell lines.

The contribution of this paper is twofold. First, we investigate whether we can predict MOA at the molecular level without optimisation of the MOA classes to the screen specificities. To this end, we benchmark a set of algorithms within a conventional pipeline, and evaluate their MOA prediction performance according to a statistically rigorous framework.

Second, we extend this conventional pipeline to the simultaneous analysis of multiple cell lines, each with potentially different morphological baselines. For this, we propose multitask autoencoders, including an adaptive model used to construct domain-invariant feature representations across cell lines. We apply these methods to a pilot screen of two triple negative breast cancer (TNBC) cell lines as models for two different molecular subtypes of the disease.

In Section 2, we describe our data set. In Section 3 we formalise a range of profiling approaches from the literature according to four key properties, and extend this to a multi-cell-line analysis. In Section 4 we illustrate the benefit of multi-task models for our dataset through extensive cross-validation and provide an exploratory analysis of differential drug effects between the two cell lines. In Section 5 we discuss our methods and the obtained results.

## 2 Data

We acquired image data for two triple-negative breast cancer (TNBC) cell lines, MDA-MB-231 and MDA-MB-468 (hereafter MDA231 and MDA468), thus constituting a multi-cell-line drug screen^1^. MDA231 (TP53, KRAS, BRAF) and MDA468 (TP53-PTEN) were both established from a pleural effusion of two different patients with triple negative metastatic breast carcinoma. Using transcriptomic and genomic data, we have recently shown that MDA468 clustered in a breast cancer specific subgroup but MDA231, one of the most used breast cancer cell lines, clustered in a mixed subgroup with cancer cell lines of very different origins, such as ovarian, urinary and kidney [30].

Both of our cell lines were subjected to the same inventory of drugs on separate 384-well microtiter plates (microplates): 36 wells contained the negative control dimethyl sulfoxide (DMSO); two the positive controls (Olaparib, Cisplatine); 166 the test compounds; and 184 empty. For each well of our two microplates, images were taken in four non-overlapping fields of view (fields), with three multiplexed fluorescent channels: (1) DAPI (cell nuclei) (2) Cyanine 3 (Cy3, *γ*H2AX to mark DNA double-strand breaks), and (3) Cyanine 5 (Cy5; tubulin marker). Together, these fluorescent channels paint a rich, composite picture of the cell populations.

The drugs comprise of a set of panels of kinase, protease and phosphatase inhibitors and can be categorised into 70 MOA classes of varying sizes, according to their targets. For our experiments, we take the 8 MOA classes having at least five member drugs. These are CDK inhibitors, cysteine protease inhibitors, EGF receptor kinase inhibitors, MMP inhibitors, DMSO (negative control), PKC inhibitors, protein tyrosine phosphatase inhibitors, and tyrosine kinase inhibitors.

In comparison with other datasets, [1] used 51 drugs in 13 MOA categories, [32] used 35 drugs in six MOA categories, and the widely studied Broad Institute Benchmark Collection 21 (BBBC21v2) [20] – used, for example, in [16] and [10] – consists of 39 drugs in 13 categories. The key difference is that our own MOA classes were not selected *a posteriori* to reflect visually different phenotypes, mounting a greater bioinformatic challenge than the standard benchmark datasets, where even a simple model can be extremely effective. For example, [31] achieved 90% accuracy with element-wise averaging of hand-crafted features after a simple luminosity correction.

## 3 Methods

This section describes the approaches for phenotypic profiling we have bench-marked. We embed these descriptions in a formalised overview of phenotypic profiling strategies to motivate the different setups. In Section 3.3 we describe methods for a joint analysis of multiple cell lines.

### 3.1 MOA prediction

Drugs are assigned a class based on their *mechanism of action* (MOA), the cellular pathway perturbed by the drug, as depicted in Figure 1. Given a set of drug profiles annotated with MOA classes, we can simulate reference and discovery drug sets in a *leave-one-compound-out* cross-validation (LOCOCV) scheme. At each fold of the cross-validation, we hold out a drug and predict its MOA class using a classifier trained on the remaining “reference” drugs. The prediction is made as the nearest neighbour (1-NN) in cosine distance between drug profiles, *d*(**p**, **p**′) = 1 − cos *θ*_**p**,**p**′_. This was proposed in [19] as an equitable way of comparing profiling algorithms. We settle for this lightweight approach as our focus here is on the discriminative power of the profiles.

**Figure 1:**
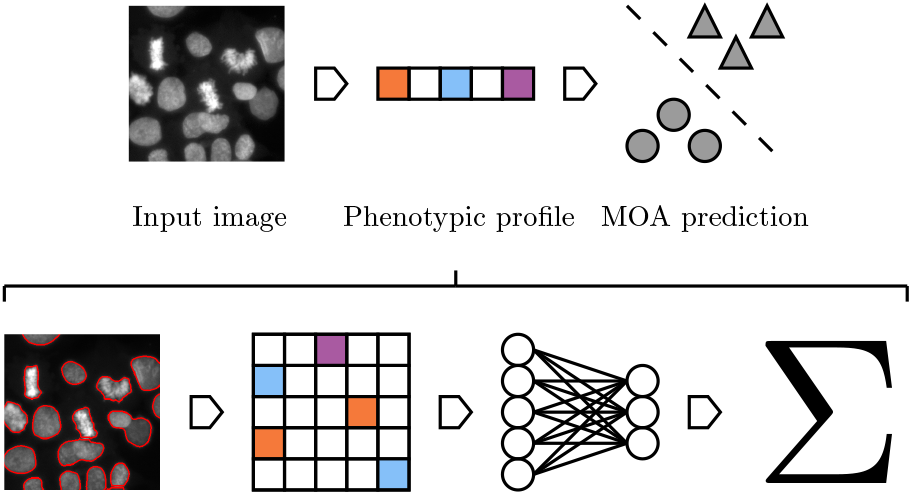
MOA prediction is performed on an image via a phenotypic profile. The development of such a profile spans four ordered stages. Each stage may be accomplished by a variety of algorithms, the combination of which define a unique pipeline. Some stages may be omitted in certain pipelines, or subsumed to a common framework.

### 3.2 Phenotypic profiling

The conventional approach to HCS analysis is a multi-stage pipeline, consisting of a sequence of modules of image and statistical analysis, including cell segmentation and hand-crafted feature extraction [6]. The aim is to ascribe a phenotypic profile to each cell population to serve as the basis of comparison between drugs. Each profile will take the form of a vector **p** ∈ ℝ^*D*^ of some dimensionality *D* and is constructed according to four ordered methodological stages: measurement unit, feature representation, dimensionality reduction, and aggregation strategy (Figure 1). Certain properties may be omitted by some approaches, or subsumed to a common framework, such as a neural network, that may perform each task simultaneously [17, 10]. In the following sections we detail each property in turn, providing references to the relevant literature and describing the concrete setup that was retained for the benchmarking.

#### 3.2.1 Measurement unit

The most common measurement unit is the cell itself, constituting a *per-cell* analysis. This entails an initial segmentation of the cells (their nuclei and other organelles). We segmented cell nuclei on the DAPI channel by subtracting a background image formed with a mean filter, before clipping to zero. Touching nuclei were further separated by applying the watershed transform on the inverse distance map of the foreground image. The cytoplasm was segmented from the microtubule channel (Cy5) following [15].

Alternatively, one might analyse the image field directly in a *per-image* analysis, such as in [25], [35], or [10]. Such approaches are referred to as *segmentation-free*, as they obviate the segmentation phase of the conventional pipeline. In this study, we deliberately choose to focus on the cell as unit of measurement.

#### 3.2.2 Feature representation

For a given choice of measurement unit, one further chooses a feature representation. This yields a matrix **X** ∈ ℝ^*N×D*^ for each well where *N* is the number of samples for that well and *D* is the number of features measured. For each segmented cell we extracted a previously published set of features [37] across the three fluorescent channels, as well as spot features informative on DNA double-strand breaks [5]. These features, hereafter referred to as *handcrafted features*, thus retain a degree of biological interpretability. In contrast, in [25] and [35] a large number of handcrafted features are extracted over each image as a whole.

More recently, features are extracted within the layers of a convolutional neural network (CNN) trained directly on image pixels. We benchmarked a convolutional autoencoder (CAE) following the design of [33], trained on 40 × 40 × 3 inputs, formed by extracting 100 × 100px padded bounding boxes of segmented cells, rescaling, and stacking the fluorescent channels. The central hidden layer of the trained CAE is then used as a feature representation.

#### 3.2.3 Dimensionality reduction

Dimensionality reduction requires some function *enc*: **X** → **Z** where **Z** ∈ ℝ^*N×M*^, with reduced dimensionality *M* < *D*. The objective is to capture the essential information in lower dimension or to cast the high-dimensional feature vector to an interpretable representation. Supervised classification of individual cells [24] is one way of achieving this, as each cell is represented either by a one-hotI:binary vector **z**_*i*_ ∈ {0, 1}^*M*^ or by a vector of probabilities **z**_*i*_ ∈ [0, 1]^*M*^ where ∑_*j*_ *z*_*ij*_ = 1 and *M* is the number of classes, in effect, the new dimensionality. With multiple-instance learning (MIL) [17] one can circumvent the manual effort involved in creating a phenotypic ontology and a manually curated training set. Here, one labels each cell with the MOA of the drug of the population, thus creating a weakly supervised ground truth. As individual cells may respond differentially to perturbation, not all regions of an image will bear the hallmarks of a particular drug, but the cellular landscape can be viewed as a multiple instance bag of objects. [10] make this assumption implicitly. We benchmarked a random forest tuned to 500 trees, trained on cells weakly labeled by MOA class of their well (*M* = 8). Necessarily, we partition wells into separate train and test sets, where the test data alone is used to build profiles for the MOA prediction downstream.

Another popular option is to use unsupervised learning. We benchmarked hard clustering methods k-means and hierarchical clustering in Euclidean space with Ward linkage. These were tuned to *M* = 80 and *M* = 100 clusters respectively (by cross-validation, on the training set). K-means is fast to fit approximately, in particular using mini-batch training. On the other hand, even using optimised software [23], hierarchical clustering is not scalable. We also performed soft clustering with Gaussian mixture models (GMM) [32], tuned to *M* = 100 Gaussians.

Feature selection [21] and principal components analysis (PCA) are other popular options. Here, we applied PCA on the handcrafted features, selecting 40 of the 516 components, retaining ~90% of the energy on average. We further whitened the latent features.

Autoencoders, as used by [16], formulate a function *f* (x) = *dec*(*enc*(x)), where *enc*(·) and *dec*(·) correspond to the encoder and decoder parts of the neural network. This model can be trained with a mean square error (MSE) loss function,

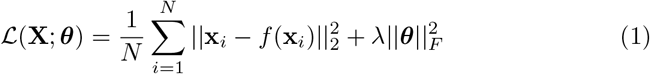

for the *N* samples in the dataset and where *λ* is a tunable hyperparameter for the regulariser. The hidden representation corresponds to the output of the encoder, the central layer of the neural network, i.e. our reduced sample is z_*i*_ = *enc*(x_*i*_). We train shallow affine autoencoders–with a single hidden layer (tuned to *M* = 100 neurons)–on handcrafted features. We also train deep convolutional autoencoders directly on image pixels, as described in Section 3.2.2. Note that such models perform both feature extraction and dimensionality reduction simultaneously. Here, the encoder consists of 5 × 5 and 3 × 3 convolutional layers, with 16 and 8 kernels respectively, and a fully connected layer (*M* = 128), each alternating with max pooling layers. The decoder mirrors this, albeit replacing pooling with upsampling.

#### 3.2.4 Aggregation strategy

Once all cells are endowed with a representation, one needs some means of reducing the population to a single profile, **p**. A variable number of cells per well requires an aggregation strategy yielding a profile of fixed size. The most straight-forward approach is an element-wise averaging as in [1] where 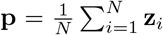. This amounts to replacing the cell population cluster with its own centroid, and for classification or clustering approaches (Section 3.2.3), this simply corresponds to the percentage of cells that fall into each category.

Alternatively, [28] apply an element-wise Kolmogorov-Smirnov test and [21] use the vector normal to the SVM decision boundary between perturbed and control populations. As we are modeling the negative control as one of our ground truth classes, we aggregate exclusively with element-wise averaging in our analysis.

### 3.3 Multi-cell-line analysis

One can extend the above MOA prediction framework for multiple cell lines either by pooling data or by ensembling models. In a pooling analysis such as [38], the cells of the respective cell lines are first normalised and then grouped across drugs to increase the amount of available data. An ensemble approach such as in [29] creates models for each cell line and aggregates their individual predictions. This approach has the additional advantage of allowing different imaging modalities of fluorescent markers.

We adopted a pooling approach to predict MOA from multiple cell lines. The challenge of this approach is to reconcile the inherent differences between the cell lines in feature space, which derives from the fundamental morphological differences of the cell lines. For this purpose, we tested multi-task autoencoders (Figure 2), extensions of both our affine and convolutional autoencoders.

**Figure 2:**
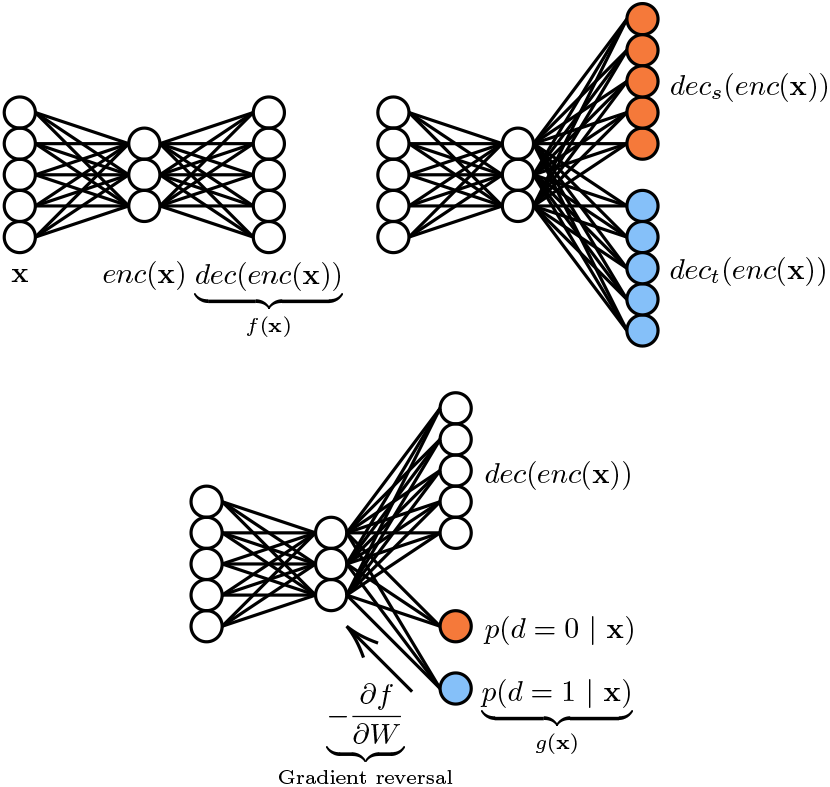
Multitask autoencoders used for dimensionality reduction over multi-cell-line data. Clockwise from top left: vanilla autoencoder, multitask autoencoder, and domain-adversarial autoencoder. Colouring indicates separate treatment of each domain (cell line).

#### 3.3.1 Multitask autoencoders for multi-cell-line analysis

Multi-task models learn to predict multiple targets simultaneously and multitask neural nets often build more generalised internal representations [7]. We propose multitask autoencoders as an approach to reconcile the divergent nature of our multi-cell-line data.

One obvious design is to have separate decoders for each cell line with a shared encoder. During training, minibatches can be split after the shared layers with samples routed to the decoder corresponding to their cell line. We thus minimise,

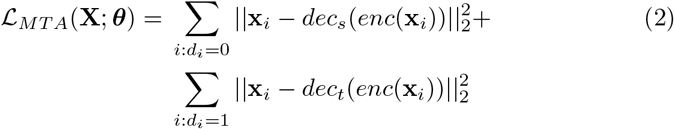

where *d*_*i*_ identifies the cell line of x_*i*_. We test multitask variants of both our affine and convolutional autoencoders described in Section 3.2.3.

The fundamental morphological differences between the cell lines can be quantified in feature space by a 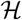-divergence, first proposed by [4], where 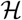 is some hypothesis class (such as the space of linea classifiers . This is expressed as 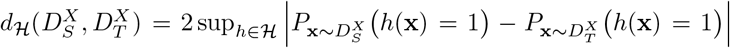, where the domains 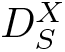 and 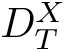 are marginal probability distributions on x. That is, given source and target domains, and given a hypothesis class 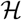, the divergence between the source and target domains is the best performance among that class of classifiers trained to distinguish them. In practice, we can approximate this by training a classifier of the class 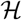 on the constructed dataset, *U* = {(x, 0): x ∈ *S*} ⋃ {(x, 1): x ∈ *T*}, that is, a classifier trained to distinguish between the domains. [2] proposed multi-task classifiers involving a domain discriminator trained against a classifier adversarially. As the classifier was trained to minimise one loss, the competing domain discriminator was trained to maximise another loss, such that data from either domain could not be distinguished, promoting *domain-invariant features* in the earlier, shared layers of the network.

Thus, we propose domain-adversarial autoencoders (DAA), to promote domain-invariant representations between the cell lines. This consists of attaching a domain discriminator *g*(x) to the encoding layer. This can be thought of as a dynamic regularisation function. In a bias-variance tradeoff, we expect this to on average *increase* the reconstruction error of the autoencoder. However, we hypothesise that the domain-invariant features learned will be more useful to the downstream MOA prediction when combining heterogeneous cell line data. For example, with a single additional affine layer, 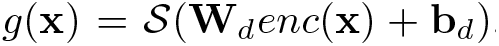, where **W**_*d*_ and b_*d*_ are the weights and biases of the layer, and 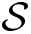 is the softmax function producing posterior probabilities *p*(*d* = 0|x) and *p*(*d* = 1|x). The loss function then becomes,

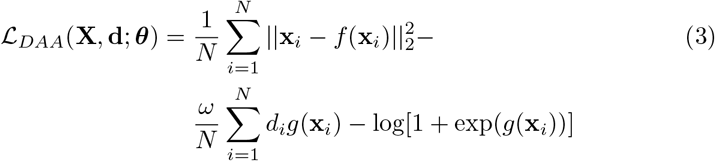

that is, the difference of a mean square error (MSE) loss and a log loss, where *f*(x) is defined as before, and *ω* is a modulating hyperparameter. However, now the parameters of *g*(x) are updated to *maximise* 𝓛_*DAA*_, so as to improve domain discrimination. At the same time, the parameters of *f*(x) are updated to *minimise* 𝓛_*DAA*_. This has the dual effect of minimising the MSE (as usual) but also maximising the log loss. This is known as an adversarial step, and aims at converging to a saddle point between the two objectives. In practice, this is implemented with a *gradient reversal* pseudo-layer [9], which is readily programmable in standard deep learning frameworks.

We test multitask versions of both our affine and convolutional autoencoders, and compare them directly in Section 4.2. The domain discriminator of our DAAs are linear in terms of the encoding (domain invariant features) and the weight of the log loss was tuned to *ω* = 1.5. For each affine model we tried the same range of hidden units in a grid search *M* ∈ {100, 125, 150, 175, 200}, and trained for 20 epochs using the RMSprop gradient descent algorithm [34]. For the convolutional autoencoders we kept the architecture defined in Section We further used weight decay (*λ* = 10^−3^) for all models as well as batch normalisation [14], which we found stabilised the training, in particular the adversarial training.

### 3.4 Model evaluation

We compared different profiling settings by evaluating performance on a MOA prediction task. For this, we balanced our datasets by randomly sampling five drugs from each of the 8 classes specified in Section 2, analysing 40 drugs at a time. Applying the LOCOCV scheme described in Section 3.1, we note that random accuracy is 12.5%. To account for random variability, we repeated LOCOCV 60 times with different sets of randomly sampled drugs and in Section 4 report average top-1 accuracy and standard deviation as the percentage of MOAs correctly predicted by the 1-NN classifier. We consider this to be a more rigorous approach in a comparative study, as while a given method often fit one drug set well, it was harder to find hyperparameter choices that worked well across all sets. We used a Wilcoxon signed-rank test to establish significance against baselines over the 60 rounds.

### 3.5 Software

We use Cell Cognition [12] to perform the first stages of the classical analysis pipeline, namely, image preprocessing, cell segmentation and feature extraction.

All models were coded using the `scikit-learn` [26] and `Keras` [8] frameworks for Python, unless otherwise noted^2^. Basic image processing was performed with `scikit-image` [36].

## 4 Results

In Section 4.1 we evaluate a range of approaches to dimensionality reduction–as described in Section 3.2.3–on their utility in creating cell representations that aggregate into discriminative phenotypic profiles for MOA prediction. This we do in separate single-cell-line experiments. In Section 4.2.1 we show how our best performing model on single-cell-line data–the autoencoder–may be extended for multi-cell-line analysis, providing comparisons for learning on handcrafted features, as well as raw pixels. We then illustrate how our optimised phenotypic profile design can be used to identify differential drug effects between cell lines across our entire drug panel in Section 4.2.2. In Section 4.2.3 we explore how the effect of adding cell lines to an analysis effects MOA predictability.

### 4.1 Single cell line analysis

In Table 1 we evaluate a range of approaches to dimensionality reduction on cell lines taken separately. The baseline for this comparison are the hand-crafted features averaged element-wise from segmented cells in each well. Note that even such a simple baseline proved to be highly competitive in earlier comparative studies such as [19]. The models are used to create a reduced representation of cells prior to aggregation by element-wise averaging (Section 3.2.4). The one exception is the convolutional autoencoder, which learns cell representations directly from image pixels.

**Table 1:**
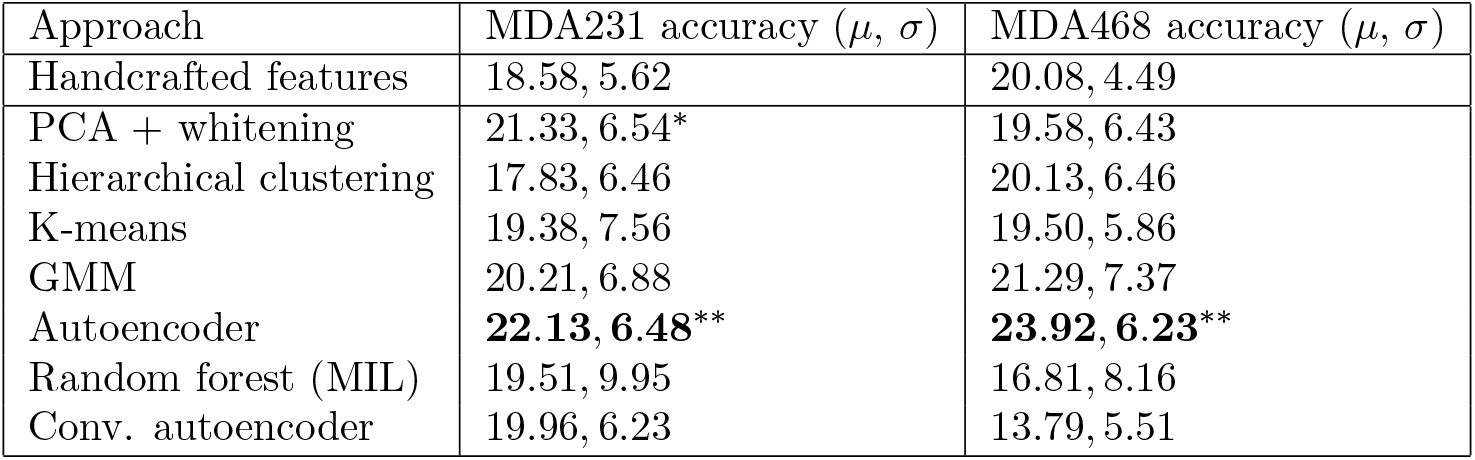
Comparison of dimensionality reduction approaches against unreduced baseline for cell lines treated separately. We show mean and standard deviation of accuracies over 60 runs with (*) indicating significant results at the *p* = 0.05 level; (**) at the *p* = 0.01 level.

| Approach | MDA231 accuracy ($\mu, \sigma$) | MDA468 accuracy ($\mu, \sigma$) |
| --- | --- | --- |
| Handcrafted features | 18.58, 5.62 | 20.08, 4.49 |
| PCA + whitening | 21.33, 6.54* | 19.58, 6.43 |
| Hierarchical clustering | 17.83, 6.46 | 20.13, 6.46 |
| K-means | 19.38, 7.56 | 19.50, 5.86 |
| GMM | 20.21, 6.88 | 21.29, 7.37 |
| Autoencoder | **22.13, 6.48\*\*** | **23.92, 6.23\*\*** |
| Random forest (MIL) | 19.51, 9.95 | 16.81, 8.16 |
| Conv. autoencoder | 19.96, 6.23 | 13.79, 5.51 |

We observe dimensionality reduction techniques register broad improvement over the baseline, with PCA (*W* = 385.0, *p* < 0.05) and autoencoders (*W* = 281.5, *p* < 0.01) significant for the MDA231 cell line. Autoencoders also registered significant improvement (*W* = 356.0, *p* < 0.01) for the MDA468 cell line. This further motivates autoencoders as the benchmark in our multi-cell-line analysis (Section 4.2).

The deep convolutional autoencoder fails to stand out from the group. However, this may rather testify to the effectiveness of handcrafted features on cell line data–at least at this resolution–over learning representations from scratch.

The sole weakly supervised method, multiple instance learning (MIL) with random forests, shows promise on cell line MDA321, but falls short on MDA468. This may stem from the necessary splitting of data into train and test sets prior to LOCOCV, reducing the available training data. Approaches based on weakly supervised MIL are popular, particularly for deep learning approaches, but we do not see any benefit for them on our dataset.

### 4.2 Analysis on multiple cell lines

So far, we have considered the analysis of several cell lines as independent problems to inform model selection. We now turn to a joint analysis on multiple cell lines.

#### 4.2.1 Prediction of MOA from multiple cell lines

With their different transcriptional programs multiple cell lines potentially bear complementary information on the mechanism of action of a drug. We pool cells in corresponding wells across our two cell lines, thus enlarging the available data for each drug. In each case, the data from each cell line were standardised separately to have zero mean and unit variance for all features. Our multitask autoencoders are compared with their single-task counterparts, the best performing models from Section 4.1.

We observe in both Tables 2 and 3 that we obtain a higher degree of accuracy in MOA prediction for our multitask autoencoders compared with their baselines, particularly the domain-adversarial autoencoders, which achieve a statistically superior average accuracy (*W* = 283.5, *p* < 0.01) for the shallow variant, based on handcrafted features, as well as for the deep learning variant (*W* = 438.5, *p* < 0.01). The former constitutes our best overall accuracy in MOA prediction on this dataset. This supports our hypothesis that promoting domain invariant features facilitates the mixing of heterogeneous data from multiple cell lines. As anticipated in Section 3.3.1, adversarial training did not improve the reconstruction error of our autoencoders, but the resultant features performed better downstream in the MOA prediction pipeline.

**Table 2:** MOA prediction on multiple cell lines (pooled) with autoencoders trained on handcrafted features. From top to bottom: vanilla autoencoders (baseline), multitask autoencoders and domain-adversarial autoencoders. We compare with the vanilla autoencoder (top row) ((**): *p* < 0.01)

| Approach | Pooled cell line accuracy ($\mu, \sigma$) |
| --- | --- |
| Autoencoder | 31.67, 6.43 |
| Multitask autoencoder | 32.04, 6.88 |
| Domain-adversarial autoencoder | **35.67, 6.94\*\*** |

**Table 3:** MOA prediction on multiple cell lines (pooled) with convolutional autoencoders. From top to bottom: vanilla convolutional autoencoders (baseline), multitask convolutional autoencoders and domain-adversarial convolutional autoencoders. We compare with the vanilla convolutional autoencoder (top row) ((**): *p* < 0.01).

| Approach | Pooled cell line accuracy ($\mu, \sigma$) |
| --- | --- |
| Conv. autoencoder | 19.58, 6.98 |
| Multitask conv. autoencoder | 20.42, 6.14 |
| Domain-adversarial conv. autoencoder | **22.38, 5.91** ** |

Inspired by [9], we use t-SNE [22] to project a sample of learned cell features into two dimensions. We typically observe a higher degree of alignment between the feature distributions of the two domains as produced by the domain-adversarial model, as illustrated in Figure 3. To quantitatively confirm this domain overlap, we compute the mean silhouette score over all points where the cluster identity of each point is simply its domain class. The scores given in Figure 3 of 0.11 (lower overlap) and 0.01 (higher overlap) for unadapted and adapted features respectively are typical.[3] wrote that multiple modalities render aggregation over a cell population problematic, as a centroid may be a bad representative of the overall population. Computing domain invariant features appears to be a partial remedy to this when pooling heterogeneous data in a multi-cell-line analysis.

**Figure 3:**
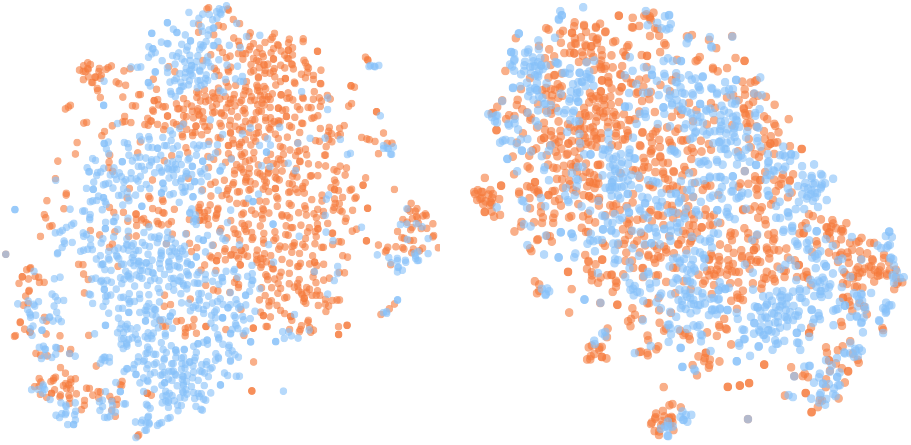
t-SNE embeddings of encodings from autoencoder (left) and domain-adversarial autoencoder (right), with cell lines distinguished by colour, and mean silhouette scores of 0.11 and 0.01 respectively.

#### 4.2.2 Differential drug effects across cell lines

Our DAA approach provides us with a representation that is optimised with respect to both MOA prediction accuracy and domain invariance between the cell two lines. This can assist us in producing profiles for all drugs in our pilot screen and investigate the differential effects of drugs across cell lines. For this, we trained our network on all data, producing phenotypic profiles for all drugs in the screen. We zero-centered each cell line by subtraction of their respective DMSO centroids and compared distances of drug profiles both from the DMSO centroid and between cell lines. By ranking these distances, we can identify four drug effect cases:

- no drug effect in either cell line;
- drug effect in one cell line only;
- differentiated drug effects in both cell lines;
- similar drug effects in both cell lines.

We visualise the relative distances between drug profiles using multi-dimensional scaling (MDS) on Euclidean distance in Figure 4 and identify examples of each of these cases. We include a comparison of DMSO populations that illustrate the unperturbed morphological differences between the two cell lines. We first show an indicative sample of DMSO cells from each cell line. Among the drugs, Endothall has a phenotypic effect on MDA231, but no visible effect on MDA468 (MDA231 cells are rounded up and smaller than in DMSO). Conversely, CL-82198 has an effect on MDA468 cells (cells are smaller and display cytoskeletal changes) and no visual effect on MDA231 cells. Cyclosporin A has a similar effect on both cell lines; the cell lines actually preserve many of their morphological baseline differences, but have a higher fraction of binucleated cells. PKC-412 has a differential effect on both cell lines. While the cell size is increased, the morphological properties as well as the number of DSBs seem to be very different between cell lines. See Supplementary Figure 1 for each drug visualised separately.

**Figure 4:**
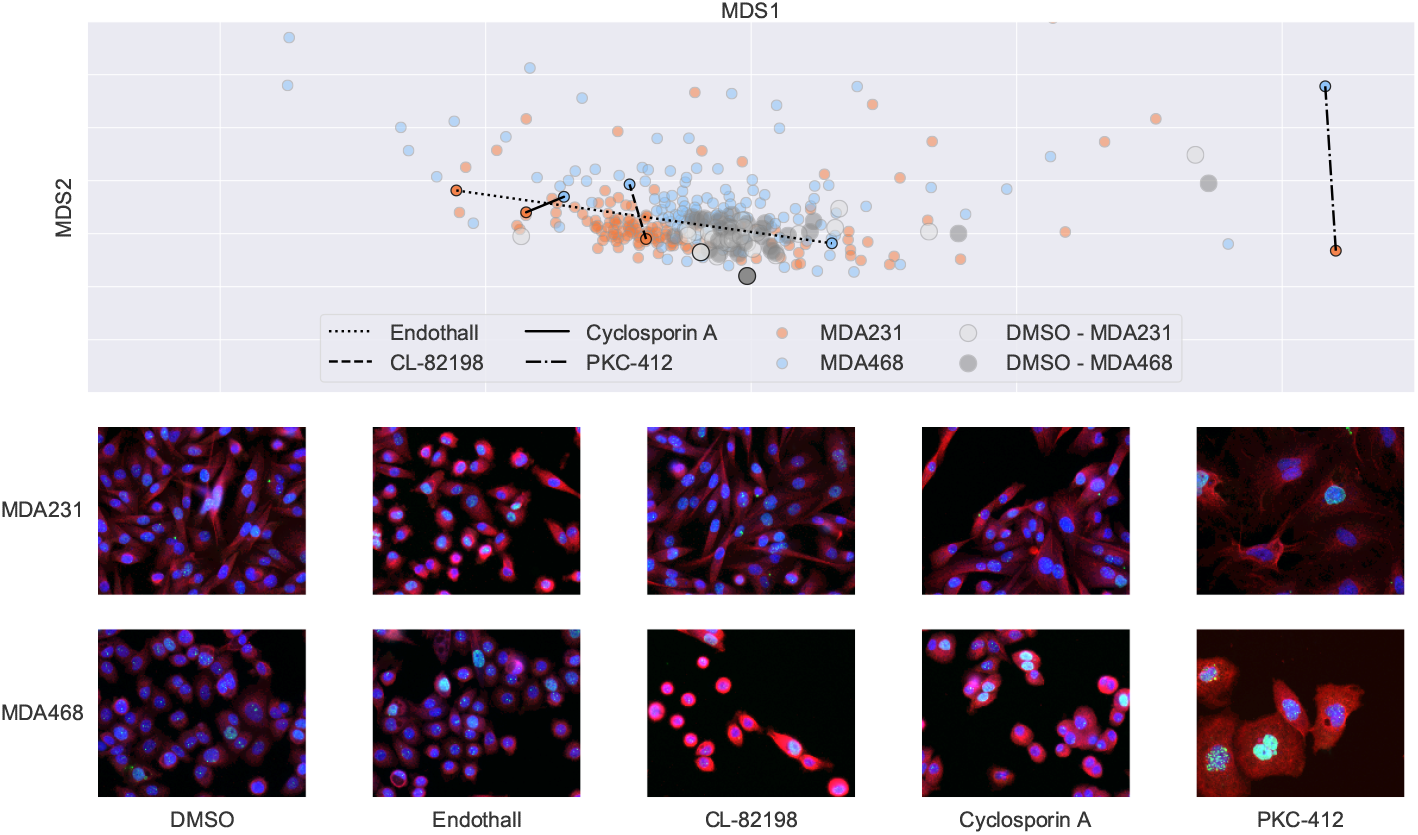
MDS embedding of drug effect profiles for MDA231 and MDA468 cell lines with DMSO centroid centered on origin. Detection of differential drug effects between cell lines with examples for each category below (MDA231 top, MDA468 bottom). From left to right: no drug effect in either cell line (negative control); drug effect in MDA231 cell line only; drug effect in MDA468 cell line only; similar drug effects in both cell lines; differentiated drug effects in both cell lines. Shown are example images, blue: DAPI, red: microtubules, green: DSB.

#### 4.2.3 Effects of accumulating cell lines

[29] demonstrated an increasing accuracy in MOA prediction as data from cell lines are added to create a growing ensemble of predictive models. This illustrates the value of drawing upon several biological sources to guide a drug discovery process. Nevertheless, predictive models will tend to perform better when supplied with greater volumes of data anyway. Any attribution of a model’s success to a richer biological foundation must first correct for the confounding effect of an increasing sample size.

We ran a separate experiment controlling for the aforementioned bias to attempt to measure the effective power of heterogeneous cell line data. To do this we created equally sized samples: 10000 randomly subsampled cells from the MDA231 cell line; 10000 randomly subsampled cells from the MDA468 cell line; and 5000 cells sampled from each cell line and pooled into a multi-cell-line dataset. We did this for the handcrafted features of segmented cells, averaged element-wise, again repeated over the 60 experimental folds. We found the pooled samples yielded an average accuracy of 20.89, significantly improving over the pure MDA231 sample at 14.94 (*W* = 240.0, *p* < 0.01) and the pure MDA468 sample at 19.42 (*W* = 516.0, *p* < 0.1). This therefore supports the hypothesis that a multi-cell-line analysis can be advantageous in and of itself, even before accounting for any increased sample size.

#### 4.2.4 Generalising to further cell lines

To test how our model behaves with increasing numbers of cell lines, we acquired image data without drug perturbation of a third TNBC cell line (MDA-MB-157) under the same protocol as the pilot screen. In order to apply domain adaptation to *K* cell lines with *K* > 2, the log loss in equation 3 was replaced with a cross entropy loss,

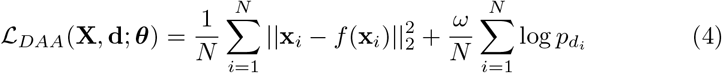

where 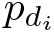 is the softmax probability produced by the domain discriminator, indexed by the domain of *i*th sample, and all other terms are defined as before. We found that this simple modification sufficed to train effectively on the new dataset, provided *ω* was reduced as the number of cell lines increased. In Supplementary Figure 2 we produce a t-SNE plot akin to Figure 3 and observe a similar tendency of distributional overlap for three cell lines. Interestingly, there is only a moderate number of additional parameters when we add a new cell line, which contrasts to the multi-task autoencoders where a whole new decoder is required for each new cell line.

As stated above, this new dataset was not a drug screen and therefore we could not evaluate our model in the same way as before. We were, however, able to pretrain our model on this morphological screen and transfer it to our drug screen to be used as a feature encoder directly. Compared with an equivalent vanilla autoencoder our model performed marginally better for three cell lines (32.08 compared with 31.17) (*W* = 366, *p* < 0.1), following our evaluation strategy. Though not the exact intended application of our model, we again see an improvement over baselines, suggesting an aptitude of our method for analysing multi-cell-line data. It will be the subject of future work to test our method on a full drug screen in greater numbers of cell lines.

## 5 Discussion

In this paper we address prediction of mechanism of action (MOA) at a molecular level. Importantly, we have not optimised the MOA classes with respect to the readout of the screen, as is common in many benchmarking studies. We have studied a number of different approaches, including traditional approaches based on hand-crafted features, and deep learning approaches, allowing us to learn suitable representations.

A major gain can be achieved by using multiple cell lines, but the choice of algorithm is important to most benefit from the data heterogeneity. We investigated several approaches, and obtained the best results for an autoencoder with a domain discriminative component to promote domain-invariant features across multiple cell lines. This approach requires the same set of markers to be used and ideally the same set of drugs to be tested.

In addition of improving MOA prediction accuracy, this method further produces a representation that allows us to compare effects of drugs on different cell lines. We use the representation in order to make comparisons between (drug, cell line) pairs. This is one of the most important use cases if the cell lines represent different molecular subtypes of a disease. Importantly, it allows one to identify highly specific drugs that only act on one particular subtype–the paradigm of precision medicine–and to distinguish them from drugs that are generally effective across different subtypes, as well as from drugs that lead to different phenotypic effects, which in turn suggest a target of different pathways depending on the transcriptional program.

While these approaches have only been applied to a small-scale pilot study, they provide an interesting starting point for larger multi-cell-line screens.

## Footnotes

^1^The full image set for this study is available at https://zenodo.org/record/2677923

^2^Worked examples of code and feature data available at https://github.com/jcboyd/multi-cell-line

