## Supplementary Figure 1 for "Domain-invariant features for mechanism of action prediction in a multi-cell-line drug screen"

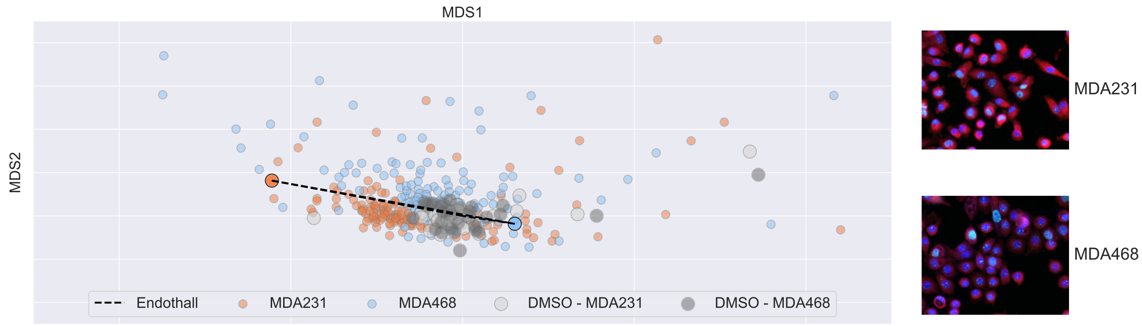


(A) Endothall takes effect in cell line MDA231 only.

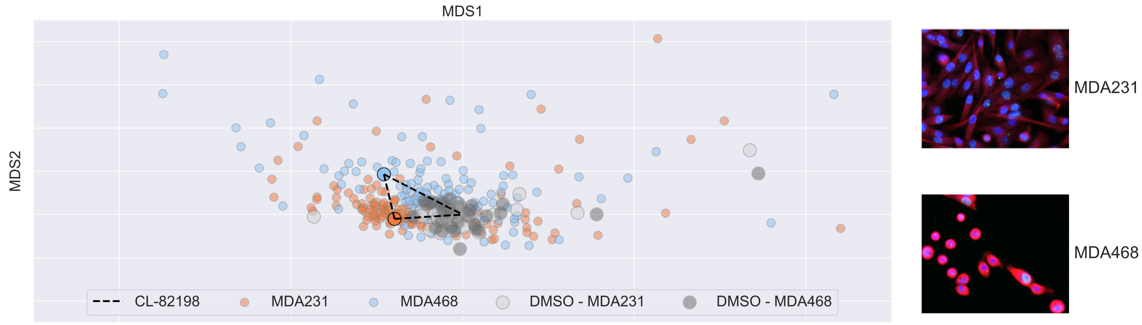


(B) CL-82198 takes effect in cell line MDA468 only.

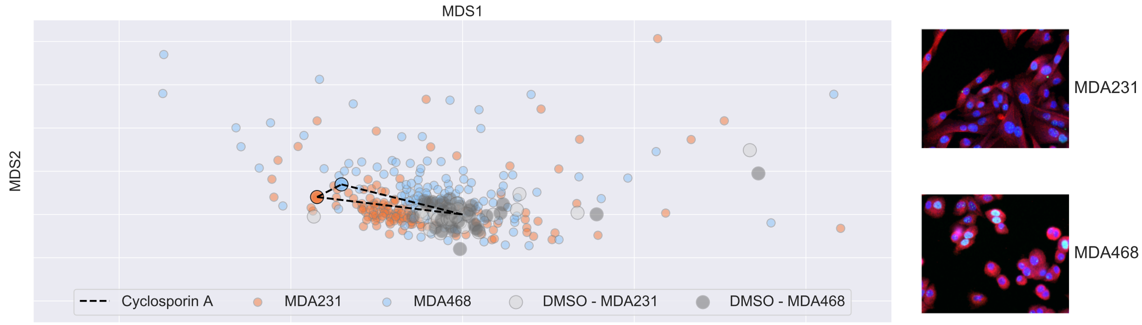


(C) Cyclosporin A takes a similar in both cell lines.

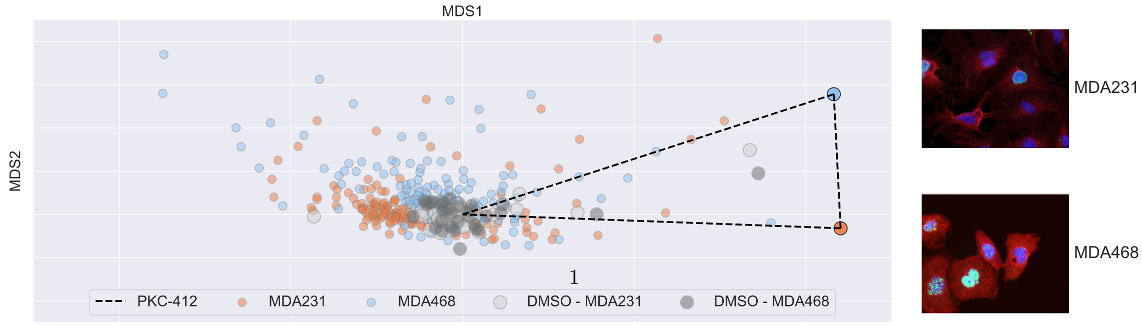




(D) PKC-412 takes differential effects in the two cell lines.

FIGURE 1. MDS plots of each category of drug effect. The distances between the profiles are plotted as a line, as well as the respective distances to the centroid (origin).
