## Supplementary Figure 1 for "Domain-invariant features for mechanism of action prediction in a multi-cell-line drug screen"

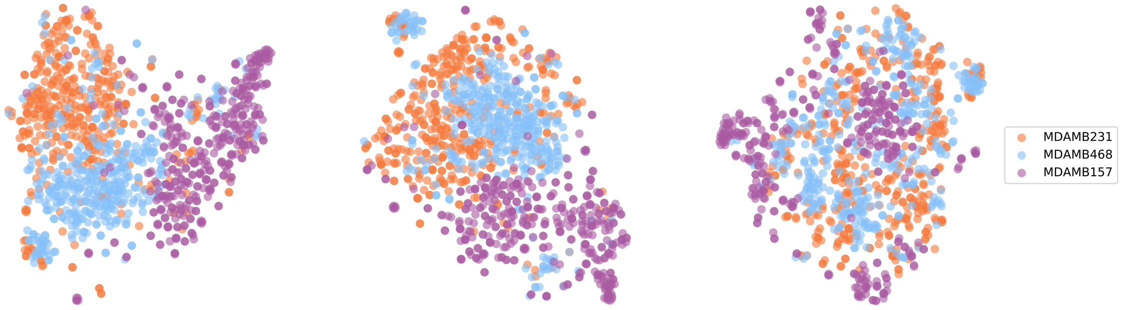


FIGURE 1. t-SNE embeddings of encodings from handcrafted features (left), autoencoder (center) and domain-adversarial autoencoder (right), with cell lines distinguished by colour. Respective silhouette scores of 0.22 and 0.14 and $-0.02$ confirm the reduced divergence in the adapted domains.
